# Reliable imputation of spatial transcriptome with uncertainty estimation and spatial regularization

**DOI:** 10.1101/2023.01.20.524992

**Authors:** Chen Qiao, Yuanhua Huang

## Abstract

Imputation of missing features in spatial transcriptomics is urgently demanded due to technology limitations, while most existing computational methods suffer from moderate accuracy and cannot estimate the reliability of the imputation. To fill the research gaps, we introduce a computational model, TransImp, that imputes the missing feature modality in spatial transcriptomics by mapping it from single-cell reference. Uniquely, we derived a set of attributes that can accurately predict imputation uncertainty, hence enabling us to select reliably imputed genes. Also, we introduced a spatial auto-correlation metric as a regularization to avoid overestimating spatial patterns. Multiple datasets from various platforms have demonstrated that our approach significantly improves the reliability of downstream analyses in detecting spatial variable genes and interacting ligand-receptor pairs. Therefore, TransImp offers a way towards a reliable spatial analysis of missing features for both matched and unseen modalities, e.g., nascent RNAs.

## 1 Introduction

A variety of biological processes are modulated through the spatial organization of cells, including how different cell types are distributed in a microenvironment and how cells communicate and perform a cooperative biological function. Prominent examples include localization of cell types in mouse organogenesis [1] and human thymus development [2], as well as inter-cellular communications in squamous cell carcinoma [3] and during intestinal development [4].

In recent years, the rapid development of spatial transcriptomics (ST) technologies has made it further accessible for dissecting the spatial mixture of cells in a wide range of biomedical research. The main two streams of technologies are sequencing-based and imaging-based (via in situ hybridization or in situ sequencing) [5]. The former in principle can cover the whole transcriptome but has a limited resolution of cells (e.g., around 5-10 cells per spot) while the latter can have a cell-level resolution but is generally limited to probing dozens of pre-selected genes [6]. Recently, breakthroughs on both platforms, e.g., seqFISH+ [7] and Stereo-seq [8], are addressing these limitations in different aspects, while the RNA capture efficiency is still far from perfect in sequencing-based methods and laborious designing of the candidate gene probes is required in the imaging-based methods.

Therefore, computational methods for feature imputation are highly demanded in analyzing ST data, particularly by leveraging the rich single-cell RNA-seq data as a reference, including imputing unseen genes in the imaging-based data or imputing lowly covered genes in sequencing data. In general, modality integration methods can be applied for the task of missing feature imputation, e.g., Liger [9] and Seurat v3 [10]. Recently, multiple tailored methods have also been proposed to address this challenge with improved performance reported. For example, SpaGE was introduced to address this task by averaging k-nearest neighbors (kNN) from the reference scRNA-seq data after projecting both ST and scRNA-seq datasets into a common low dimensional space which is spanned by adapted principal vectors [11]. Similar kNN-based aggregation strategies were also applied in a joint representation space produced by either shared principal components [1] or the latent encodings of an auto-encoder [12]. Tangram was another appealing method that directly learns a mapping matrix for cells from scRNA-seq to spots in ST by minimizing the cosine distances at both feature and sample levels [13] between imputed and true ST expressions. Moreover, in addition to certain molecular features, mapping meta information of cells, e.g., cell type labels, is also a task that shares the same principles of feature imputation but is usually treated as a standalone task, e.g., in RCTD [14] and Cell2location [15].

However, multiple challenges in ST imputation remain less touched. First, there is no indicator available for assessing the imputation reliability: it is not clear how reliable one imputed gene could be used for further biological discovery. Second, most of the feature imputation methods do not explicitly consider the spatial pattern strengths during the imputation, often resulting in overestimating spatial smoothness. Third, given the rapidly increasing number of cells in the ST data, computational efficiency is another demanding property.

To address these challenges, we introduce a generic framework, TransImp, to transform information from a scRNA-seq reference to the ST context, with two major innovations. First, it can provide uncertainty scores for imputation performance, hence allowing us to focus on genes with more confident imputation. Second, it introduces a regularizer for spatial pattern preservation, alleviating the over-estimate of spatial auto-correlation. To demonstrate the effectiveness of our model, we focused on a few challenging tasks, including the prediction of the dominant proportion of missing features in the image-based ST datasets. We also verified its high reliability in common downstream spatial analyses: detection of spatially variable genes and interacting ligand-receptor pairs. Finally, we briefly showcase that this method can also be applied to predict the unspliced RNAs, hence enabling trajectory analysis of cell differentiation in a physical space.

## 2 Results

### 2.1 TransImp imputation and uncertainty inference

In TransImp model, we aim to learn a mapping (i.e., translation) function *f*(·) to translate the scRNA-seq reference to ST data. Related to the Tangram model [13], our overall translation framework is to learn a linear mapping matrix *W* from *N*_*c*_ reference cells to *N*_*s*_ ST spots (Fig. 1a). One can also view it as a multivariate regression model (i.e., multiple outcome variables), by treating genes as samples and cells as feature dimensions (see the difference of this dual problem in Supp. Fig. S1). Here, we further simplified the translation function to be a linear model without bias and introduced two modes: full and low-rank (see details in Methods). To ensure computational efficiency and model robustness, we only use the low-rank mode for single-cell reference (referred to as TransImpLR or simply TransImp) and full mode for the cell cluster reference (referred to as cluster mode or TransImpCls). One may see that the cluster mode (TransImpCls) is a special case of the low-rank model by pre-defining the cell loading matrix **V** as the cell type identity matrix.

**Figure 1:**
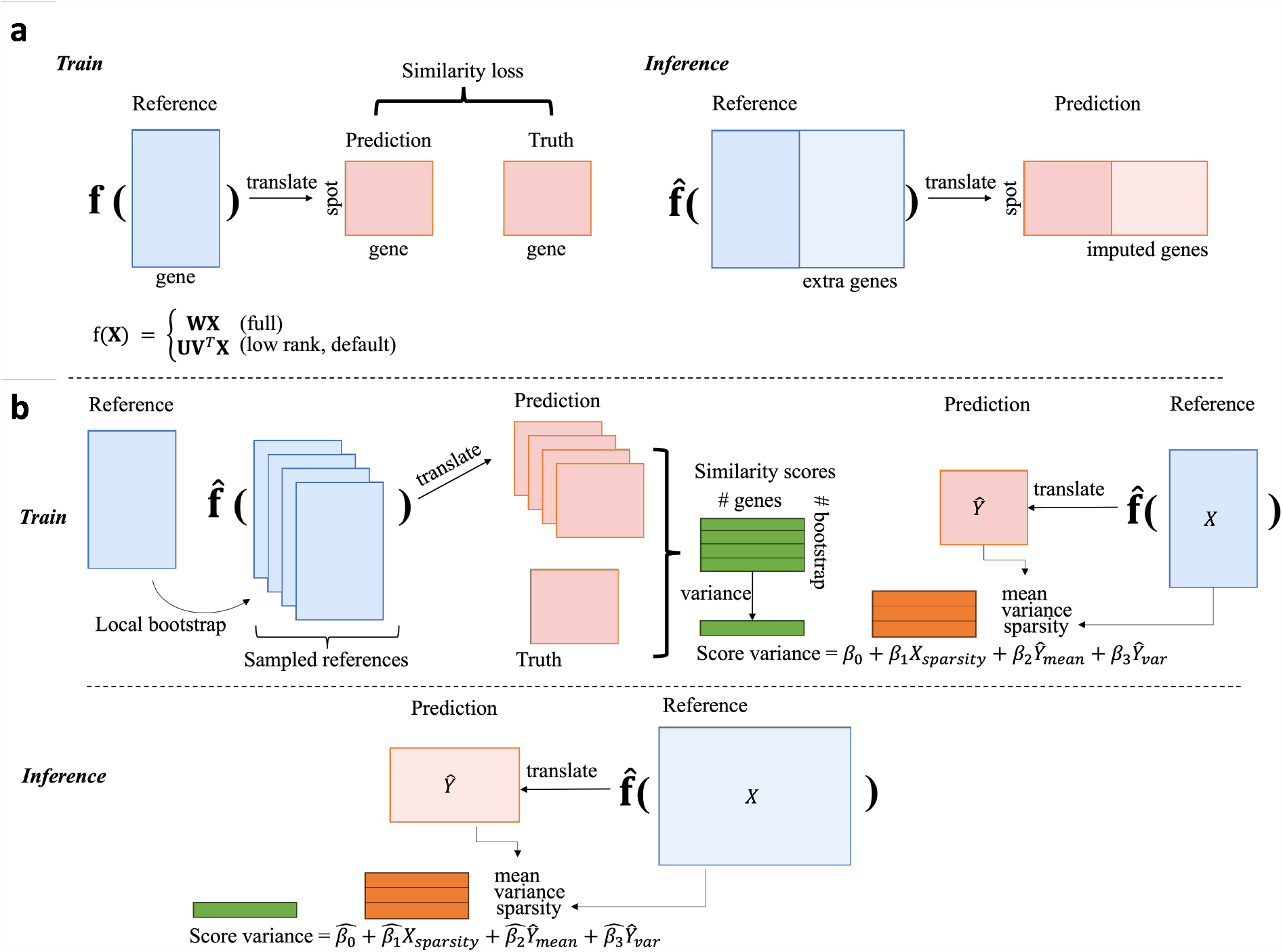
TransImp computational framework. a. The training and inference procedures of the imputation model; b. The training and inference procedures of the post-hoc uncertainty prediction model.

Then, the translation function is trained on the overlapped genes between reference and spatial datasets by minimizing the cosine similarity loss between the predicted and observed spatial expression matrices at both gene and spot levels. Once the translation function 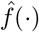 is learned, it can be applied to impute those genes that are unseen in the ST data but observed in single-cell reference data (Fig. 1a right panel). Of note, this framework can be easily extended for adding regularization terms into the loss function, for example, a spatial regularization term based on spatial auto-correlation statistics, Moran’s *I*, which we will discuss in Section 2.4 and show how it encourages the mapping function to preserve spatial patterns in translation.

The translation function alone, however, lacks an indicator of prediction confidence on those missing genes. Therefore, we propose a framework to estimate the prediction uncertainty as illustrated in Fig. 1b. First, we estimated the variance of imputation performance (Score variance) on the training genes (where we have the true ST expression) as a *post-hoc* step relying on a fitted translation function, and the same training SC reference and ST datasets. Specifically, given the estimated translation function 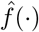, sampled reference matrices from local bootstrapping (see Section Methods) on the observed reference count matrix can all be translated into the ST domain, where similarity scores can be computed against the observed ST count matrix (Truth) for each sampling iteration. Hence, the variance of the cosine scores among all bootstrapping samples can be calculated and serve as a score variance. Second, by using those training genes, we aimed to predict the score variance of the missing/imputed genes with a linear regression. The model consumes three features for each gene: sparsity of gene reads from the reference count matrix denoted as *X*_*sparsity*_, mean and variance of the imputation prediction *Ŷ* denoted as *Ŷ*_*mean*_ and *Ŷ*_*var*_ respectively. Last, with the estimated uncertainty prediction model parameterized as 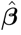, a gene’s performance uncertainty can be inferred by feeding the corresponding features of the observed reference and imputed ST expressions.

### 2.2 TransImp contributes to state-of-the-art imputation and its estimated uncertainty prioritizes unprobed ST genes for reliable analysis

We first applied TransImp to a dataset generated with the seqFISH platform on mouse organogenesis, where 351 genes were probed in 57,536 spots [1], covering 24 major cell types and their distributions as shown in Fig. 2a. To assess the imputation performance, we conducted a 5-fold cross-validation on these 351 genes and merged all the test folds for evaluation. In Fig. 2b and c, we show example genes that are well and poorly imputed, respectively. The well-imputed genes tend to better capture the ground-truth spatial patterns, while those poorly imputed genes with either spurious or weak ground-truth spatial patterns challenge the model in prediction. An overall performance comparison for different imputation methods is visualized in Fig. 2d, where the Cosine similarity score (CSS) indicates that the proposed method achieved the best performance regarding imputation consistency with ground-truth (median CSS= 0.499, 0.483 for TransImpLR and TransImpCls respectively), outperforming the existing methods at least moderately when using the same train-test split. We noticed that stPlus and Tangram also work comparably well (median CSS=0.463 and 0.477, respectively), while SpaGE is less accurate on average (median CSS=0.454).

**Figure 2:**
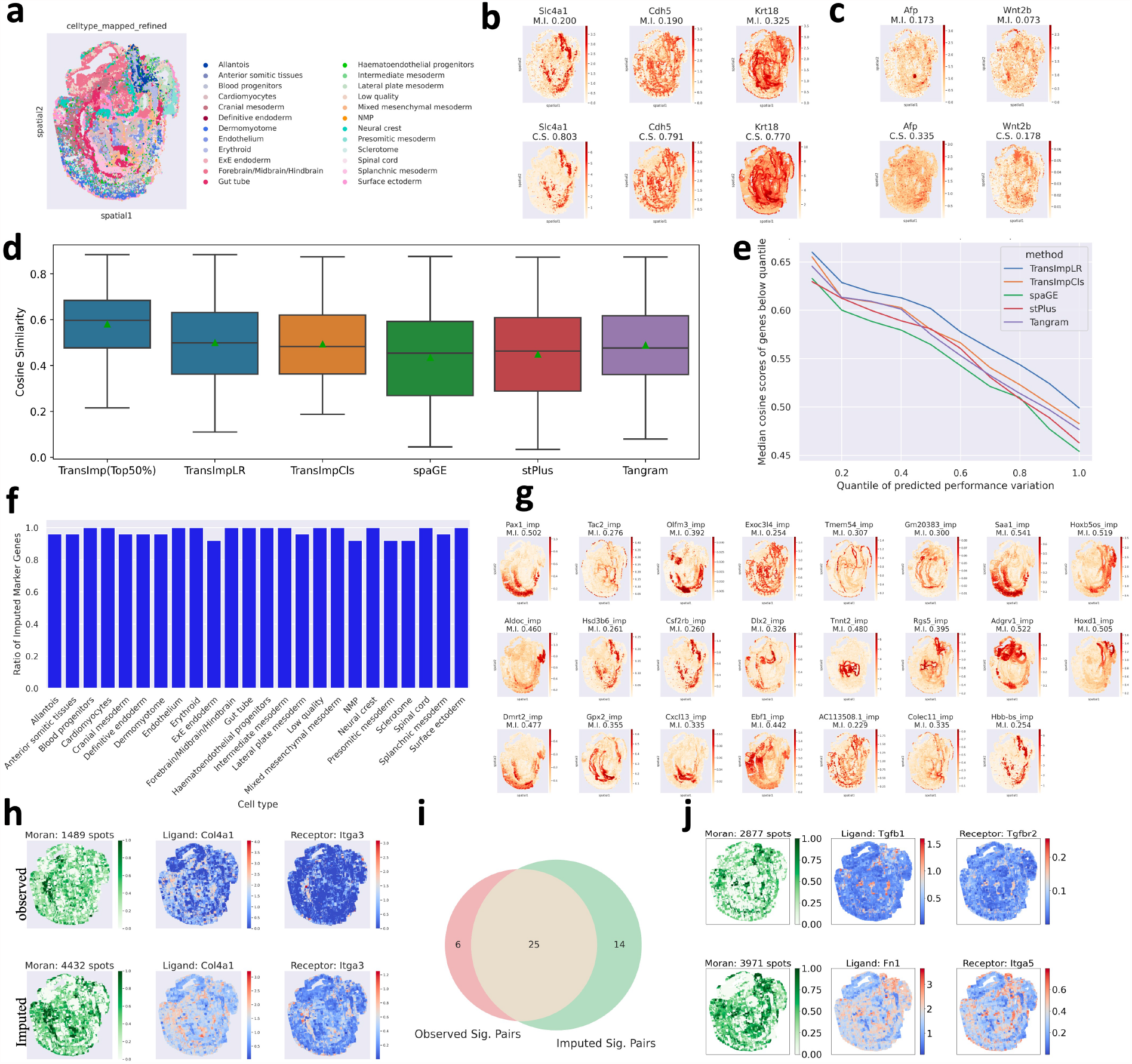
Evaluation results on SeqFISH dataset. a. Observed cell type distributions over spatial locations; b. Example well-predicted genes, upper: observed, lower: imputed; M.I: Moran’s *I* statistics; C.S.: cosine similarity score; c. Example less well-predicted genes, upper: observed, lower: imputed; d. box plots of cosine similarity scores (CSS) for all methods, the first box shows the statistics of a subset of genes with predicted uncertainty below the median of all genes. e. Line plot of CSS aggregated in different quantile ranges, the x-axis denotes the quantiles of predicted uncertainty, the y-axis denotes the median CSS of genes below the corresponding uncertainty quantile; f. Bar plot for proportions of unprobed marker genes in each cell type; g. The top-1 ranked marker genes for each cell type, postfix “img” indicates unprobed genes imputed from single-cell reference. h. Spatial pattern plots of an example spatial ligand-receptor interaction pair, upper: on observed data, lower: on imputed data. In each row, the leftmost plot shows the probabilities of spatial interaction over spots, while the rest two show the expression patterns of the involved genes. i. Vann diagram of significant ligand-receptor pairs in observed and imputed ST expression matrices; j. Plots of two spatial ligand-receptor pairs (upper and lower) of unprobed genes.

Despite achieving state-of-the-art performance, the overall accuracy is still not perfect, partly due to discrepancies between the reference and ST datasets and technical measurement noise. Therefore, we asked if our proposed performance uncertainty surrogate can help identify more confident genes (see Methods). After ranking genes by their performance uncertainty, we found that the median CSS can be substantially improved from 0.499 to 0.600 by focusing on the half gene set with lower uncertainty (Fig. 2d). Moreover, this trend is highly consistent where the median CSS of genes at lower uncertainty quantiles tend to be higher (Fig. 2e), and it is not only for our method but also for all other methods, suggesting our proposed performance uncertainty is an effective indicator of imputation confidence.

Given the enhanced accuracy of our uncertainty-aware imputation, we further explored to which extent the imputed genes will facilitate the biological analysis, including the cell type marker genes and spatial ligand-receptor interaction. When examining the marker genes from the pool of observed and imputed genes, we found >90% of markers are imputed genes for all cell types (Fig. 2f). The top-1 ranked marker genes for each cell type are shown in Fig. 2g, which turns out to be unprobed genes for all cell types. These results indicate that much richer information may be entailed in the unprobed genes, and that imputation is one solution to the limitations of ST technologies.

We further investigated gene interactions over spatial locations and used SpatialDM [16] for detecting significant spatial ligand-receptor interactions. We first run the test on all probed genes, and Fig. 2h shows a typical example pair that is significant in both observed and imputed expressions. The figure also shows that the spatial interaction patterns in imputed expressions are more widespread (4,432 vs 1,489 significant spots with local communication). When assessing all ligand-receptor pairs, 57 pairs are covered in the 351 genes, among which 31 and 39 were identified as interacting pairs by using the observed and imputed expression, respectively (FRD<0.1). In Fig. 2i, the Vann diagram further indicates a big overlap of significant pairs between observed and imputed data, implying the accuracy of imputation. Moreover, when leveraging the power of imputation on all unprobed genes, more significant interactions can be discovered (45 pairs), such as the two example pairs shown in Fig. 2j, where active interaction regions cover cell types such as Allantois and Neural crest. The results again indicate the potential values of unprobed genes for biological discovery that can be achieved via imputation.

### 2.3 TransImp is efficient and robust across datasets from multiple platforms

Next, we conducted evaluation experiments on three more spatial transcriptomic datasets generated using different technologies. In Fig. 3a, we summarized the CSS of gene profiles across datasets and methods in box plots. It is evident that our methods TransImpLR/TransImpCls consistently achieved the best performance compared to those state-of-the-art methods in cosine similarity scores. Moreover, the predicted uncertainty did well in identifying reliably imputed genes, as demonstrated in the line plots of Fig. 3b, where genes with more certain performance at the lower quantiles tend to achieve higher CSS scores. When selecting the 50% genes with lower predicted uncertainty, we can find in Fig. 3a (the first boxes) that they have much higher median CSS scores than CSS of all the genes (e.g., 0.718 vs0.562 on MERFISH data).

**Figure 3:**
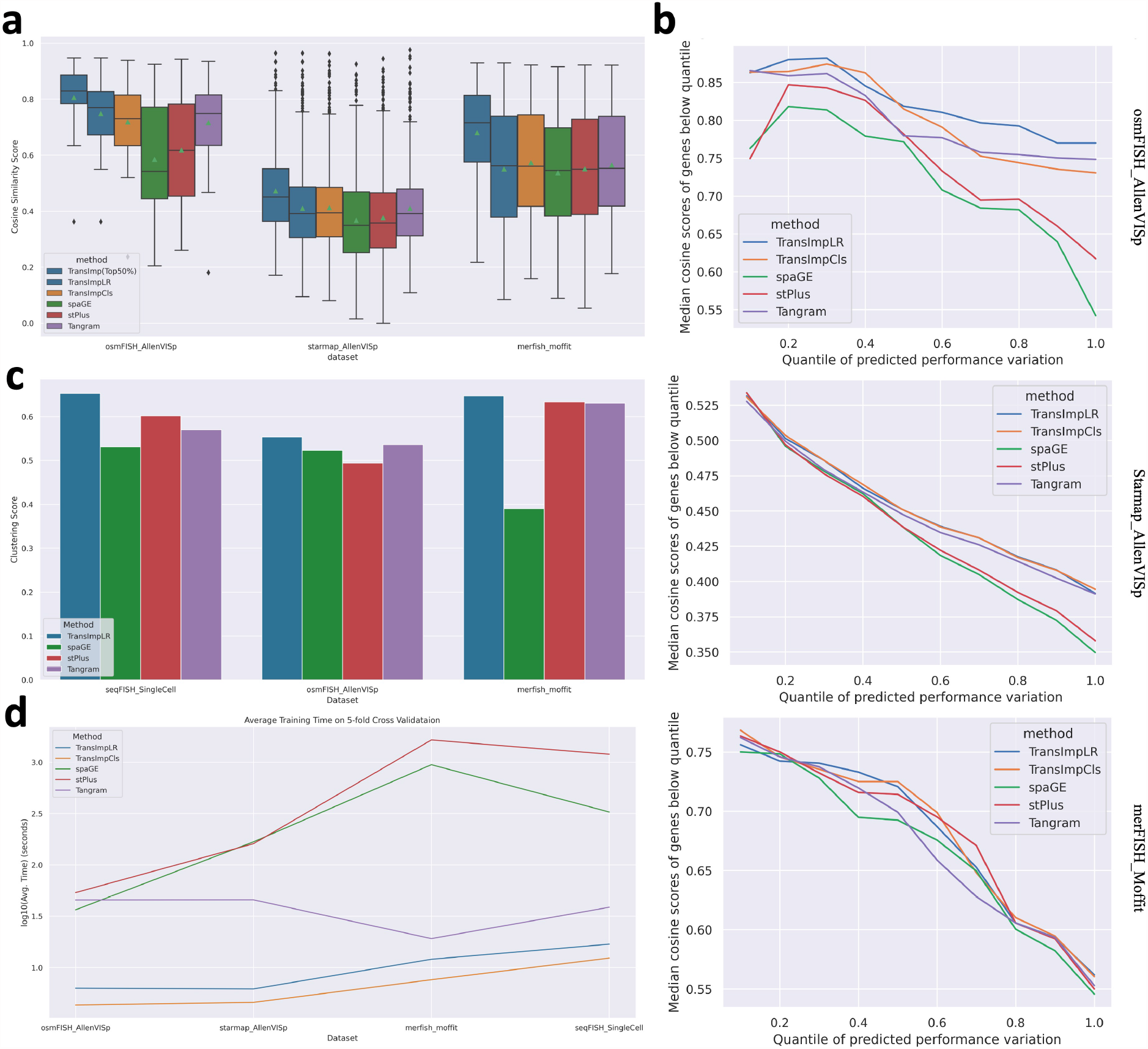
Evaluation of imputation methods. a. Cosine similarity scores (CSS) for osmFISH, starmap and MERFISH spatial transcriptomic datasets; b. Line plots of imputation performances in different uncertainty quantile ranges, where the x-axis denotes the quantiles of predicted uncertainty and the y-axis denotes the median cosine similarity score of genes below the corresponding uncertainty quantile; c. Averaged clustering scores from multiple metrics (Methods); d. Averaged computational runtime of different methods.

Moreover, downstream clustering analysis was conducted on the imputed genes from the 5-fold test sets. To involve spatial information in the clustering, we adopt Agglomerative Clustering with adjacency matrices calculated using the spatial coordinates. We compare the clusters of imputed versus true expressions after applying the same Agglomerative Clustering procedure. The averaged clustering metrics (covering multiple scores, e.g., Adjusted Rand Index; see Methods) are visualized as Fig. 3c, demonstrating our method TransImpLR consistently achieved the best performance across all three datasets, particularly with clear gain on the seqFish dataset (0.653 vs 0.602 as the second best).

Finally, we recorded the training run-time for each method. As shown in Fig. 3d, benefited from GPU acceleration, Tangram, and our proposed methods are much more efficient than stPlus and SpaGE, particularly on larger datasets such as MERFISH and seqFISH. Between Tangram and our method, we found both TransImpLR and TransImpCls can still achieve 37.1-90.5% running time reduction, probably thanks to the low-rank setting.

### 2.4 Spatial regularizer preserves spatial auto-correlation, reinforcing the downstream signal detection

Although TransImp and other methods allow the imputation of missing genes, in empirical analyses we constantly find a common interesting phenomenon: the spatial patterns of the imputed gene expressions tend to be overestimated and hence exhibit stronger Moran’s *I* statistics than observed. Taking TransImpLR on seqFISH dataset as an example, the imputation methods increase the spatial auto-correlation Moran’s *I* index (M.I) from observed to imputed expressions on the test set (Fig. 4a top two rows). Interestingly, this trend is also true if we treated the imputation as clean ground truth and added synthetic noise to make a new training set, where we found again the imputed data achieved much higher Moran’s *I* statistics (Fig. 4a bottom two rows) than the noised observation.

**Figure 4:**
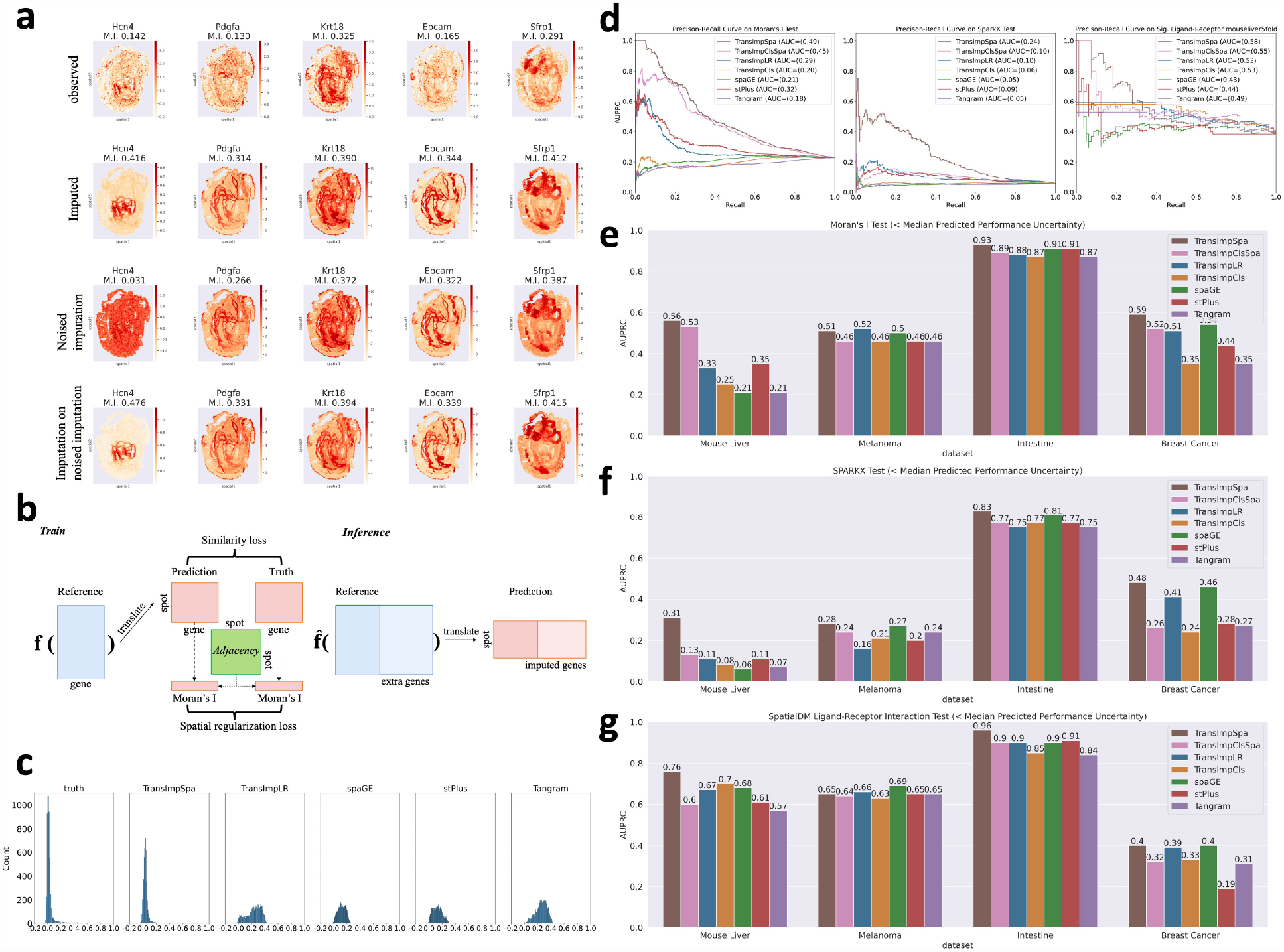
Spatialy regularized imputation and experimental results. a. Example genes from the simulation experiment for demonstrating the denoising property of imputation. Top row: Observed gene expression; second row: Imputed gene expressions that tend to have stronger moran’s *I*-detected patterns; third row: white-noised version of imputed genes from the above row, with spatial patterns weakened; bottom row: imputation results with the noised genes as training data, the spatial patterns are much higher than the noised observations; b. The training and inference procedures of the spatially-regularized imputation; c. Histogram plots of Moran’s *I*s on mouse liver dataset, indicating that the spatial regularization enables the predictions to have more consistent Moran’s *I* patterns; d. From left to right, precision-recall curves and their AUC scores of Moran’s *I* Spatially-highly variable gene (SHVG) detection tests, of Spark-X SHVG detection tests, and of spatial ligand-receptor interaction (SLRI) detection tests on mouse liver dataset; e-f. Bar plots of area under precision-recall curve scores for Moran’s *I* SHVG tests, Spark-X SHVG tests, and SLRI tests. Shown are results derived from genes with predicted performance uncertainty below the median.

Overestimating the spatial pattern can be a risk for downstream biological analyses and an ideal model should retain spatial patterns as consistent as possible with the observation. To achieve this goal, we propose to add a spatial regularization to the training objective. As an analogy to the regularizer used in LASSO and the ridge regression model, we anticipate a spatial pattern-based regularizer may prevent the model from overestimating spatial patterns. The auxiliary training loss is illustrated in Fig. 4b. As shown in the figure, in addition to the standard similarity loss, a spatial regularization loss based on Moran’s *I* is enrolled to make *I*s of the observed and predicted expressions consistent. The strength of regularizing could be adjusted by tuning its weight (see Methods). After estimating 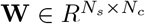, the standard inference process applies seamlessly as in Fig. 1a. To validate, we applied it to a real-world dataset and found (see Fig. 4c) other methods without spatial regularization have distribution mass on much larger Moran’s *I*s than observed ground truth (the overestimation phenomenon), while spatially-regularized TransImp, i.e., TransImpSpa, has distributions closer to ground truth.

Next, we examine whether or to what extent spatial regularization can improve the accuracy of downstream analysis for biological pattern discovery. Specifically, we applied both spatially regularized and unregularized configurations of our method (with or without the -Spa suffix) together with other methods to four Visium-based ST datasets. Thanks to Visium’s capacity to sequence almost the whole transcriptome, far more genes can be captured, and we hence can obtain enough positive and negative observations for assessing the detection of spatially highly variable gene and spatial ligand-receptor interaction. Similar to the analyses above, we used the results from observed gene expression as ground truth and examined the correctness of that from imputed expressions. On the mouse liver dataset, Fig. 4d demonstrates the better spatial pattern preservation performance of TransImpSpa with the auxiliary regularization. Overall, the tasks are extremely challenging for all the methods, yet TransImpSpa continues to obtain the best results. Specifically, for spatially highly variable gene detection (Fig. 4d left and middle), TranImpSpa achieved the highest area under the precision-recall curve (AUPRC) with a testing metric by using either Moran’s *I*(AUPRC: 0.49 by ours vs 0.32 by others) or Spark-X (AUPRC: 0.24 vs 0.09), corroborating its effectiveness of retaining spatial patterns for individual genes. Moreover, for spatial ligand-receptor interaction detection using SpatialDM (Fig. 4d right), TransImpSpa also outperforms all the other methods by a large margin (AUPRC: 0.58 vs 0.49), showing its good spatial-pattern-preserving performance even in the inter-gene interaction context.

Besides the mouse liver datasets, we further examined the robustness of the spatial regularization on three human datasets: melanoma, intestine, and breast cancer. The overall AUPRC performance scores for all the methods are shown in Fig. 4e-g for Moran’s *I*, Spark-X, and Ligand-receptor interaction tests, respectively. Of note, we used predicted uncertainty to filter out genes above median uncertainty, since Visium ST datasets are of greatly high dimension and low quality. It is noticeable from the bar charts that TransImpSpa robustly outperforms other methods without spatial regularization in finding SHVG genes with either Moran’s *I* or Spark-X methods, and it also remains the robust high-performing method for detecting spatial ligand-receptor interactions.

### 2.5 TransImp facilitates spatial RNA velocity analysis by imputing unspliced and spliced RNAs in spots

Finally, as a generic translation framework, TransImp may have the capability of translating the unseen feature modality in the ST data from the reference scRNA-seq. Here, we specifically assess how the unspliced RNA (and the spliced RNA) abundance can be translated from scRNA-seq data to ST data. To explore, we trained the TransImpSpa framework on the Chicken Heart and Mouse Brain datasets with corresponding reference datasets retrieved from [17]. On each dataset, the model was trained on the anchor genes (shared between scRNA-seq and ST) and then used to predict the unspliced and spliced expression matrices into the spatial space, followed by RNA velocity analysis with scVelo stochastic mode [18].

On the Mouse Brain dataset, we first performed the clustering on the scRNA-seq data and translated the cell type to the ST data (consistently between Tangram). As there lacks ground truth of the unspliced and spliced RNA abundance, we evaluated the performance by comparing the consistency of the predicted differentiation directional between ST and scRNA-seq. In Fig. 5a, we found that the neuron differentiation from neuroblast cells is well captured in both scRNA-seq and ST data (Supp. Fig. S5 and S6).

**Figure 5:**
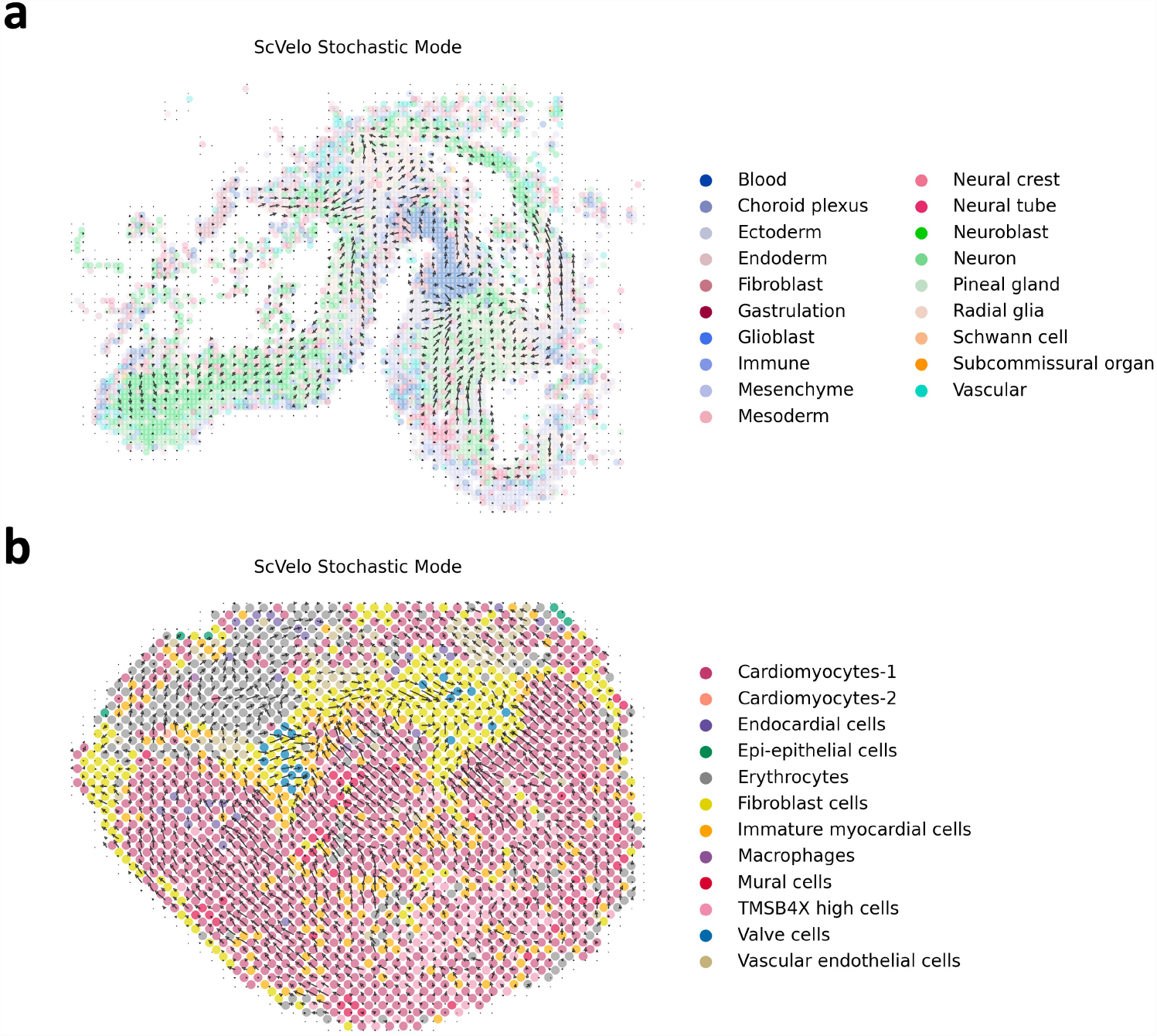
Spatial RNA velocity analysis based on imputed nascent and mature RNA counts, spot cell types are mapped from single cell annotation. a. Spot-level transition grid map on the Mouse Brain ST dataset, sub-regions of the map show transitional directions from Neuralblast to Neuron captured in single cell level; b. Spot-level transition grid map on the Chicken Heart ST dataset, the overall trends of terminating at Fibroblast cells are consistent with prior work.

By performing a similar analysis on the chicken heart dataset where the cell types were provided, we found the overall transition trends terminating at Fibroblast cells are consistent with the pseudotime analysis for the Epicardial lineage in [19]. Taken together, these results demonstrate the potential of spatial RNA Velocity analysis translating unseen modalities using the proposed framework.

## 3 Discussion

To summarize, we introduced a framework for imputing missing features from reference scRNA-seq data. The predicted performance uncertainty helps identify reliable imputations. On various datasets from different platforms, we demonstrated that the proposed framework achieves state-of-the-art prediction accuracy, the predicted genes with lower uncertainty are more reliable, and the spatial regularization preserves the spatial patterns of the imputed features. Also, the wide applicability in common downstream analyses and computational efficiency of our method may further accelerate the analysis of increasingly popular ST data.

On the other hand, it still remains an open challenge to better structure imputation models with the location information available from spatial transcriptomics. First of all, there are other model families to be explored for this task and our low-rank mapping matrix has a high flexibility to be adapted to them. As briefly mentioned, the full translation framework working with cluster-aggregated gene signature matrix (TransImpCls) can be viewed as a special case of the low-rank framework with a cell-by-gene reference matrix, in that the low-dimensional matrix **V** is fixed to be a binary matrix of shape cell-by-cluster. Each row of the matrix is a one-hot vector turned on at the dimension corresponding to this cell’s cluster type. The low-rank setting hence offers an additional interface for injecting prior knowledge (e.g., cell types) into the translation function, either via explicit regularization on **V** and/or **U** or through a Bayesian manner. Moreover, beyond a linear setting for the low-rank framework, the property of none-linearity may also be added to the translation function via, e.g., none-linear activation functions after the dot product with **V** and/or **U**, increasing model capacity for more complex mapping.

Second, as we demonstrated, local bootstrapping allows us to estimate performance uncertainty, which not only can be predicted by empirical features but also serves as an effective indicator of the reliability of the imputation. The low accuracy in the imputation for some genes may be inevitable for (almost) all computational methods, as the key information may be missed due to the immature technology, making the selection of accurate genes a necessary step for reliable biological discovery. Considering that the number of missing genes is often large, this prioritization of more accurate genes can further push the computational methods for real applications, even though we are still at the beginning of assessing and predicting the performance uncertainty.

Last, for the spatial regularization module, one may further re-evaluate spatial metrics that play an important role in quantifying spatial patterns of gene expressions, based on which the discrepancies between prediction and truth can be measured. It should be noted that the properties of a spatial metric affect downstream analysis. In TransImpSpa, we leveraged Global Moran’s *I* as a proxy for quantifying spatial patterns. However, studies have argued for more powerful metrics such as SpatialDE [20] and Spark-X [21] for mining spatial patterns. Besides, applying local spatial metrics such as local Moran’s *I* and Geary’s *C* can be achieved by enabling spot-centered mini-batch optimization for translation functions, instead of fully batched training resulted from calculating global spatial metrics. This allows scaling the spatially-regularized frameworks to even larger datasets.

## 4 Methods

As shown in Fig. 1a, a standard translation framework translates an input reference gene profile or signature into the target gene profile on spots, where we treat genes rather than cells/spots as an input instance. The translation function entails the simplest configuration where it is linear without bias terms (assuming both datasets are normalized to the same scale), reducing to an alignment/mapping matrix as inferred by Tangram [13]. Nevertheless, a translation function could be more flexible as being low-rank or even non-linear. Moreover, we may add spatial regularizations in fitting translation functions, so that they could better preserve spatial patterns.

### Translation function

A translation function *f*(·) takes as input a reference gene profile **x** and outputs the target spatial profile **y**: **y** = *f*(**x**). This work considers simple settings of being full or low-rank linear mapping:

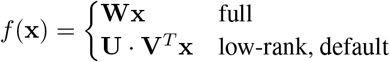

Where non-negative matrix 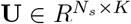, *N*_*s*_ and *N*_*c*_ are the numbers of spots and cells, respectively. In the low-rank setting, **W** is approximated by the matrix multiplication of two low-dimensional non-negative matrices 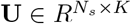 and 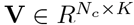, where K is a hyper-parameter specifying the dimensionality. We constrain **V** to be non-negative by taking the element-wise square of an unconstrained matrix 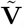, denoted as 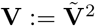, **U** is constrained by taking the softmax of an unconstrained matrix **Ũ** over the *K* latent dimensions. The design motivation is to first implicitly generate latent cluster centers (in space *R*^*K*^) and then construct spots by weighted combinations of these centers. Besides, we consider two modes of the input **x**: 1) Cluster mode, aggregating reference matrix by summing over clusters provided by e.g., Leiden method; and 2) Cell mode, the whole gene profile vector. For efficiency of computation, we apply low-rank mapping for the cell mode (as default) and full mapping for the cluster mode (a special case of the low-rank cell mode).

### Translation loss

Shared genes between reference and spatial transcriptomic datasets are used for supervised training of the translation function. We denote the output of the translation function for all the genes as matrix **Ŷ** which is compared with the ground-truth matrix **Y**. The translation loss is computed based on the cosine similarity between the rows and columns of the two matrices, capturing both the spot-wise and gene-wise expression distributions:

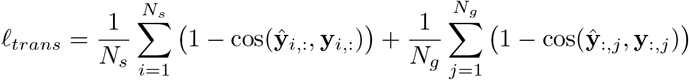

where **y**_*i*,:_ and **y**_:,*j*_ index the *i*th row and *j*th column from matrix **Y**, respectively. *N*_*g*_ is the total number of shared genes.

### Spatial regularization loss

To explicitly encode spatial patterns into the training procedure, we adopt global Moran’s *I* [22], a well-studied spatial auto-correlation metric, as the quantitative measurement, and compare *I* values on predicted and true expressions of each gene using mean squared error (MSE) loss:

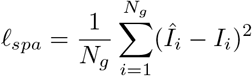

where *Î*_*i*_ and *I*_*i*_ are Moran’s I computed on predicted and true expressions of gene *i* respectively (see Fig. 4b).

### Uncertainty estimation

To estimate the reliability of gene imputation, we propose a post-hoc uncertainty prediction model as illustrated in Fig. 1b. This is a linear regression model designed to predict the uncertainty of imputation performance for each imputed gene. The dependent variable is performance uncertainty (score variance in the figure). For training data, this uncertainty score can be derived with a local bootstrapping procedure. With the single-cell reference matrix, we sample with replacement in each Leiden cluster the exact same number of cells within this cluster. After obtaining *N*_*sim*_ sampled reference matrices, the already estimated function 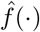 can translate all of them into the ST domain, where *N*_*sim*_ similarity scores can be computed against the observed ST expression matrix (Truth). The score variance can then be derived for each gene and serve as the training target for regression. The model consumes three features: sparsity of gene reads from the reference count matrix denoted as *X*_*sparsity*_, mean and variance of the imputation prediction *Ŷ* from the original single-cell reference, denoted as *Ŷ*_*mean*_ and *Ŷ*_*var*_ respectively. With these training data, the following linear regression model can be trained:

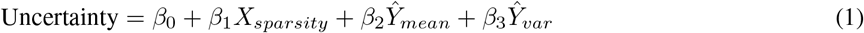

With the trained model 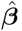, a gene’s performance uncertainty can be inferred by feeding the corresponding three features into the model. We would expect those genes with smaller uncertainty to be more reliable, by assuming that the local resampling of the original single-cell reference matrix should less affect reliably imputed genes, since the local context should be more homogeneous for well-predicted genes.

### Model configuration and training

Four settings of the proposed framework are studied and evaluated on different datasets. As shown in Table 1, configurations denoted as TransImpClsSpa and TransImpCls are Cluster-based full mapping frameworks with and without spatial regularization respectively. Likewise, TransImpSpa and TransImpLR are cell-based low-rank settings of the translation framework with and without spatial regularization.

**Table 1:** Model configurations.

|  | w/o Spa.Reg | w/ Spa.Reg |
| --- | --- | --- |
| Low-Rank (cell mode) | TransImpLR | TransImpSpa |
| Full (cluster mode) | TransImpCls | TransImpClsSpa |

For configurations with spatial regularization, a hyper-parameter *λ* is used to balance the spatial regularization strength in the total loss:

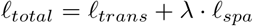

Only the translation loss *ℓ*_*trans*_ is used for configurations without spatial regularization, which can also be viewed as a special case of the total loss when *λ* = 0.

All the models are implemented using pytorch 2.0 [23] and trained with AdamW optimizer [24] on GPU.

### Datasets and configuration

Two categories of spatial transcriptomic (ST) datasets are used for evaluating the imputation performance of the proposed methods, as shown in Table 2. The top four rows summarize the imaging-based ST datasets. We obtained the preprocessed STARmap, MERFISH and OsmFISH, as well as the corresponding references AllenVISp and Moffit from [11], while the SeqFISH dataset with its single-cell reference is obtained from [1]. The bottom four rows summarize the information of the Visium-based ST datasets. The preprocessed mouse liver ST c1 sample and its reference are from [25]. The breast cancer ST sample 1142243F and single-cell reference are from [26] and [27], respectively. We achieved the preprocessed human melanoma [28] ST dataset and its reference as well as the intestine [29] ST A1 sample from the spatialDM authors [16], and the intestine reference is obtained from [30].

**Table 2:**
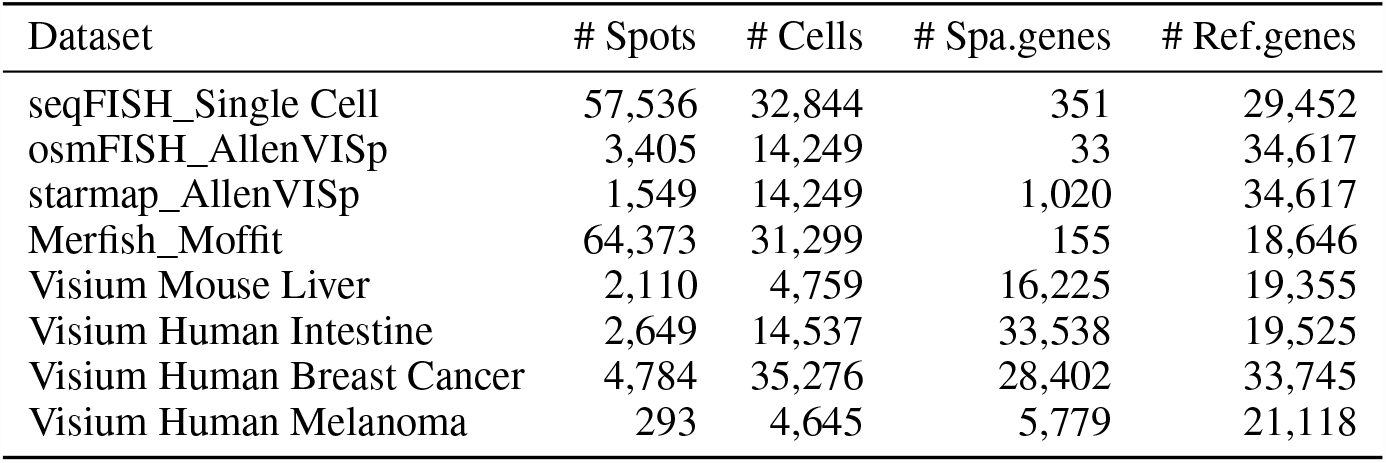
Dataset information.

For all the datasets, we train all the models with 2000 epochs, a learning rate 0.01, weight decay 0.01, and set the latent dimension for low-rank modes to be 256. There are only two exceptional TransImpSpa models on Visium datasets each requires one differently set hyperparameter: The latent dimension to be 128 on the Melanoma dataset to make it further smaller than the number of spots 293; The elements of **V** clipped to be within 0.5 for the intestine dataset to prevent overfitting.

For RNA velocity analysis, the preprocessed versions of the two spatial transcriptomic datasets with the corresponding single-cell references, Day 14 Chicken Heart [19] and Developing MouseBrain Atlas [31] are achieved from [17].

### Evaluation of imputation

We compare our method and previous methods including stPlus [12], SpaGE[11] and Tangram [13] on 5-fold cross-validataion results over different ST datasets. To measure the similarity between predicted and true gene profiles, cosine similarity score (CSS) is calculated for each gene and aggregated by median (Fig. 2d and Fig. 3a) within each dataset.

### Evaluation on spatially highly variable gene detection

To further evaluate the imputation methods, we assess the downstream task of detecting spatially highly variable genes from the imputed expression matrices. This evaluation can only be conducted on Visium-based ST datasets, due to their almost whole genome-wide sequencing capacity that can capture enough positive and negative genes for analysis. We adopted both the classical Moran’s *I* test [22] and the more recent non-parametric Spark-X test [21], and set the significance level to FDR<0.01 for both methods. Viewing significant and non-significant results as binary classification, we may draw precision-recall curves (PRC) (in Fig. 4d and Supp. Fig. S2 and S3) and summarize the performances of different methods as the area under the curve (AUC) (in Fig. 4e-f), which is a better metric than the area under the Receiver-Operating Characteristic curve in scenarios of label imbalance.

### Evaluation on spatial ligand-receptor pair detection

Beyond spatial patterns of individual genes, we evaluate methods in a more challenging task that tries to identify spatially interactive ligand-receptor pairs. The recently developed method SpatialDM [16] leverages a bivariant Moran’s statistic to detect spatial co-expression patterns of ligand and receptor pairs and is used as the assessment tool in our evaluation. We set the significance level of FDR to 0.01, and after running the test for the ground truth and all the imputed expression matrices, we could also plot PRC and calculate the summary AUC for model comparison, as shown in Fig. 4d and g, and Supp. Fig. S4.

### SeqFISH unprobed gene analysis

The single-cell reference dataset has 29,452 genes, from which we selected the top 1,000 highly variable genes and uniformly sampled 1000 genes. The intersected genes excluding those in the 351 probed SeqFISH genes amount to 1,754, which constitutes the final set of unprobed genes imputed from single-cell reference. We combine probed (observed) and unprobed (imputed) ST genes into an extended SeqFISH dataset, and ran Wilcoxon-based marker gene detection and SpatialDM implemented spatial ligand-receptor interaction detection tests.

### Spatial clustering evaluation

We further evaluated our methods on downstream clustering analysis. The analysis was conducted on OsmFISH, MERFISH, and seqFISH, where the annotation of cell types is available. Agglomerative clustering structured by a spatial adjacency matrix was applied to each dataset, including both the true and the predicted expression matrices. We then compare the clustering results of predicted and true expressions with averaged clustering indices of Adjusted Rand Score (ARS), Adjusted Mutual Information Score (AMIS), Homogeneity Score (HOMO), and Normalized Mutual Information Score (NMI) (in Fig. 3c).

### Spatial RNA velocity exploration

To explore the potential application of the proposed method on Spatial RNA velocity analysis, we fit TransImpLR on reference datasets and translate its unspliced and spliced mRNA count matrices into the spatial space, where RNA velocity was inferred using scVelo [18].

## Code availability

The TransImp is implemented as an open-source Python package and freely at https://github.com/qiaochen/tranSpa. For reproducibility, all analysis notebooks are also available in this repository with links to the pre-processed datasets.

## Data availability

Datasets preprocessed and generated in this study are published at https://doi.org/10.5281/zenodo.7556184

## Acknowledgement

We thank Miss Zhuoxuan Li for kindly sharing the pre-processed Melanoma and Intestine datasets. We acknowledge support from the University of Hong Kong and its Li Ka Shing Faculty of Medicine as well as the Centre for Translational Stem Cell Biology (CTSCB).

