## Supplemental materials for "Reliable imputation of spatial transcriptome with uncertainty estimation and spatial regularization"

### Supplementary Figures for "Reliable imputation of spatial transcriptome with uncertainty estimation and spatial regularization"

Chen Qiao and Yuanhua Huang

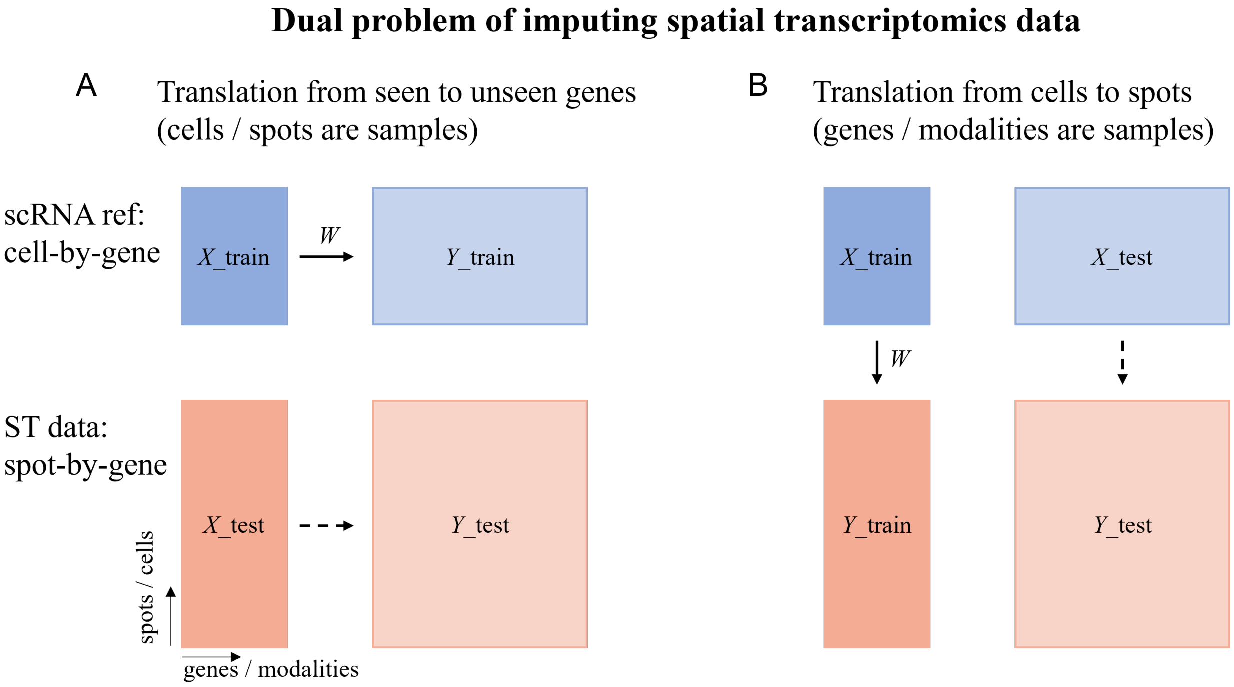


Figure S1: Dual problem of imputing spatial transcriptomics data. A. Translate observed genes to unobserved genes. B. Translate cells to spots, as used in our method.

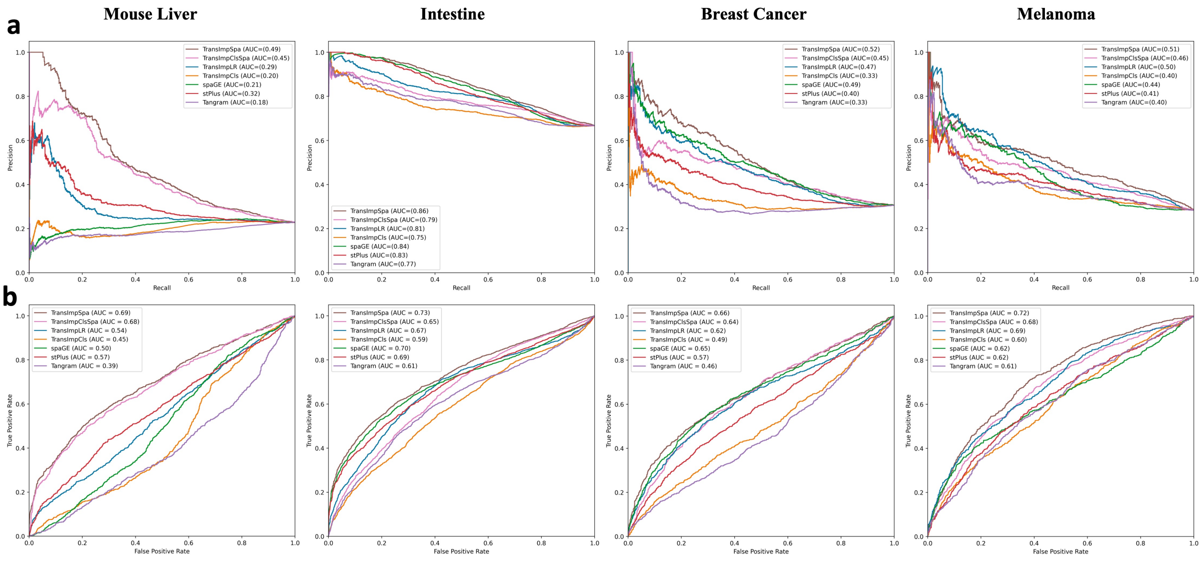


Figure S2: Results of spatially-highly variable gene detection on imputed Visium ST datasets using Moran's I test (FDR $< 0.01$). a. Precision-recall curves; b. Receiver Operating Characteristic curve.

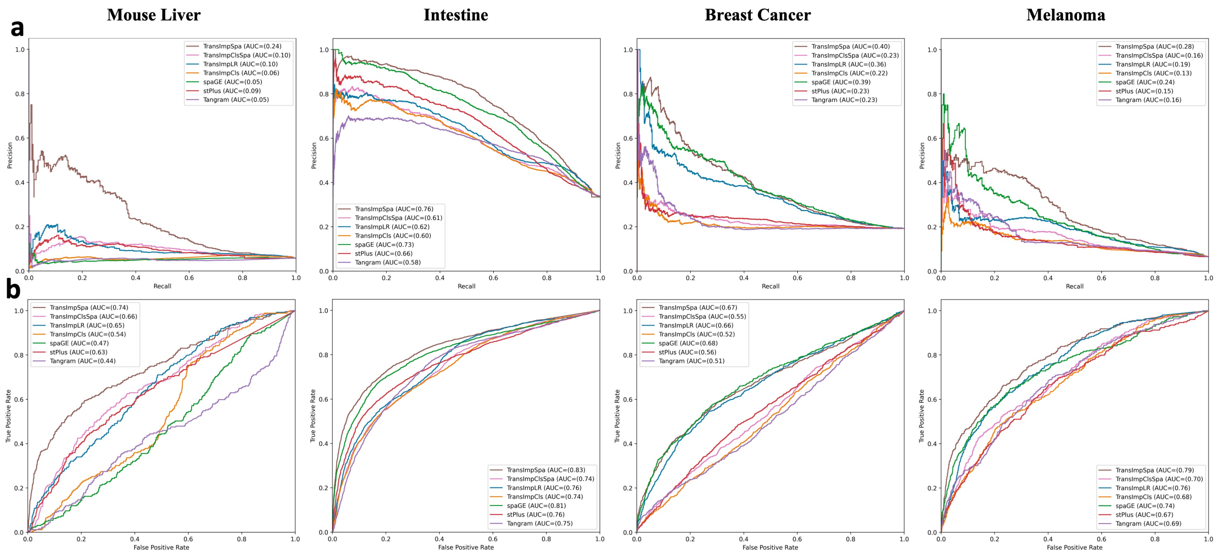


Figure S3: Results of spatially-highly variable gene detection on imputed Visium ST datasets using Spark-X test (adjusted PVal $< 0.01$). a. Precision-recall curves; b. Receiver Operating Characteristic curves.

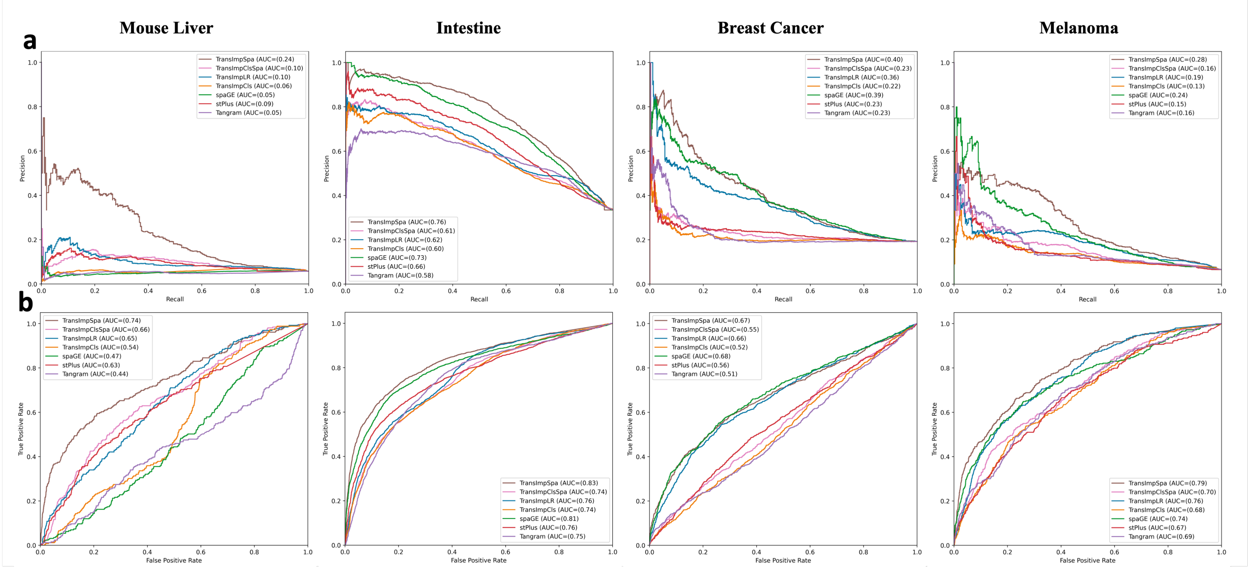


Figure S4: Results of spatial ligand-receptor pair detection on imputed Visium ST datasets using SpatialDM (FDR $< 0.01$). a. Precision-recall curves; b. Receiver Operating Characteristic curves.

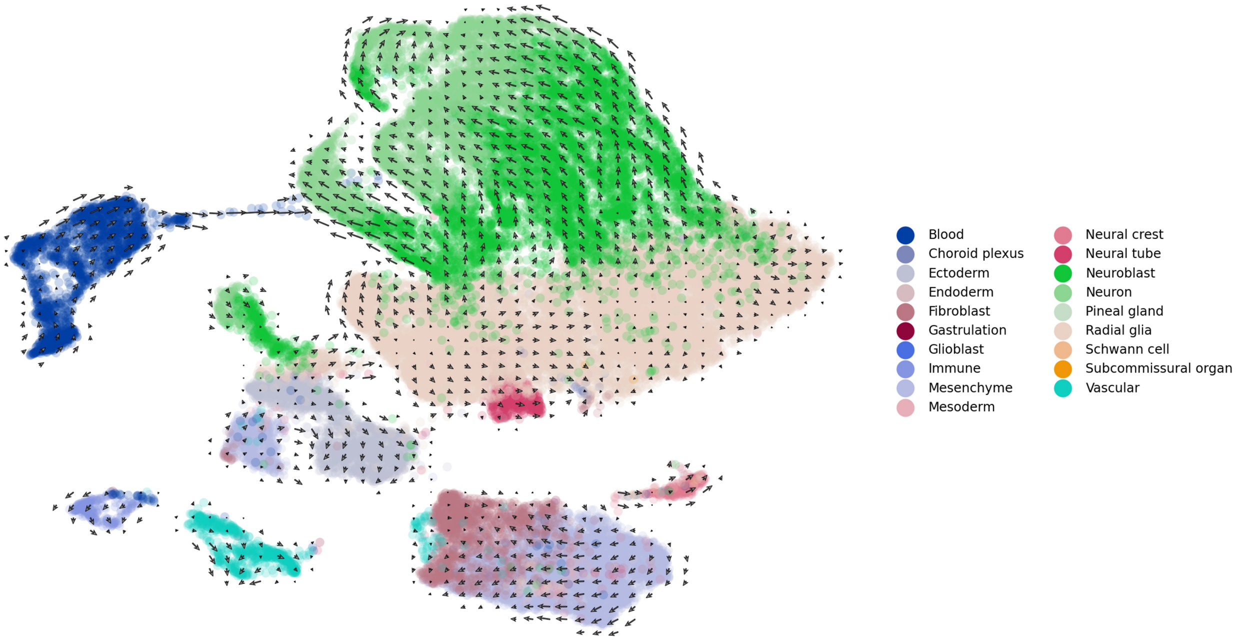


Figure S5: Cellular transition grid of the Mouse Brain data set based on RNA velocity estimated at the single cell level.

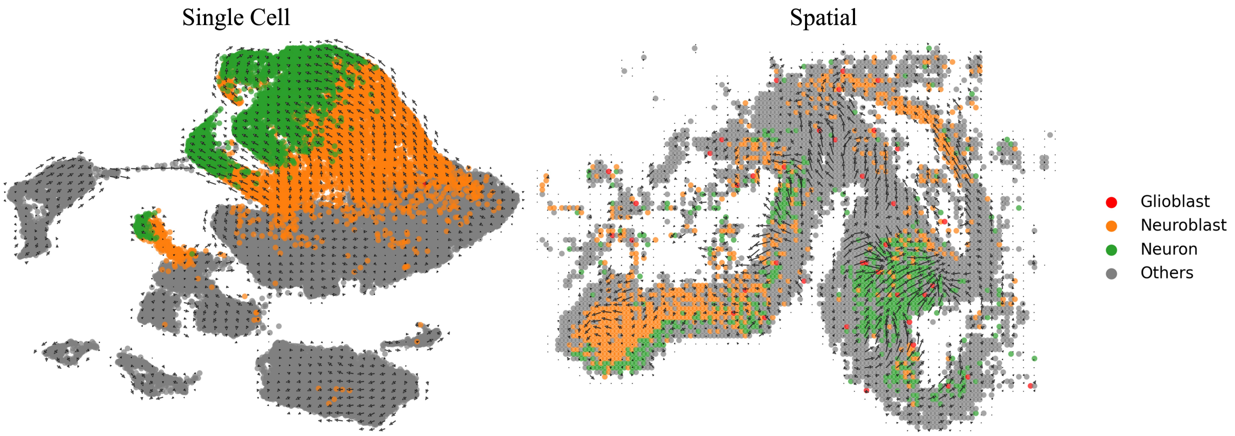


Figure S6: Cellular transition grid of the Mouse Brain at both single-cell and spatial levels colored by four cell types: Glioblast, Neuroblast, Neuron and others.
